# Untargeted metabolomics reveals a selectively enriched surface metabolome underpinning constitutive chemical defence in the red alga *Halymenia floresii* (Clemente) C.Agardh, 1817 ((Rhodophyta, Halymeniales)

**DOI:** 10.64898/2026.10.19.760206

**Authors:** Shareen A Abdul Malik, Gilles Bedoux, Yolanda Freile-Pelegrín, Nathalie Bourgougnon, Daniel Robledo

## Abstract

Marine macroalgae rely on chemically mediated surface defences to prevent microbial colonization and fouling, yet the composition of surface-associated metabolites and their ecological roles remain poorly understood. In this study, untargeted liquid chromatography–mass spectrometry (LC–MS)-based metabolomics was used to characterize the surface-associated metabolome of the tropical red macroalga *Halymenia floresii* and identify metabolites potentially involved in chemical defence. Selective dipping extraction (DIP) was used to recover surface-associated metabolites and compared with whole-cell metabolome (WCM) extracts. Across both datasets, 22,455 LC– MS features were detected, including 10,505 features in the surface extracts and 11,495 additional features detected only in the WCM extracts. Statistical filtering identified 722 LC–MS features consistently enriched by at least four-fold in surface extracts relative to WCM extracts. These features yielded 928 putative metabolite annotations using xMSannotator, with biological interpretation supported by KEGG, LIPID MAPS, and MassBank. Pathway reconstruction using the Lilikoi R package identified 44 metabolite–pathway associations, predominantly involving lipid-associated metabolism, including steroidogenesis, arachidonic acid metabolism, plasmalogen synthesis, fatty acid biosynthesis, sphingolipid metabolism and sterol biosynthesis. Manual curation further indicated enrichment of fatty acids, oxylipin-like metabolites, sphingolipids, phospholipid precursors and isoprenoid intermediates at the algal surface. Rather than indicating the presence of metabolites unique to the surface, the findings suggest that *H. floresii* selectively enriches lipid-associated metabolites at the thallus–seawater interface. Spatial localization of relatively hydrophobic metabolites may provide an energetically efficient chemical defence strategy by retaining bioactive compounds at the site where microbial attachment and biofilm formation are initiated while minimizing their loss to the surrounding seawater. Overall, this study demonstrates that chemical defence in *H. floresii* is associated with the selective surface enrichment of lipid-associated metabolites rather than the occurrence of unique surface-specific compounds, providing new insights into the spatial organization of macroalgal chemical defences and the mechanisms underlying resistance to microbial colonization and fouling.

**Graphical Abstract:** 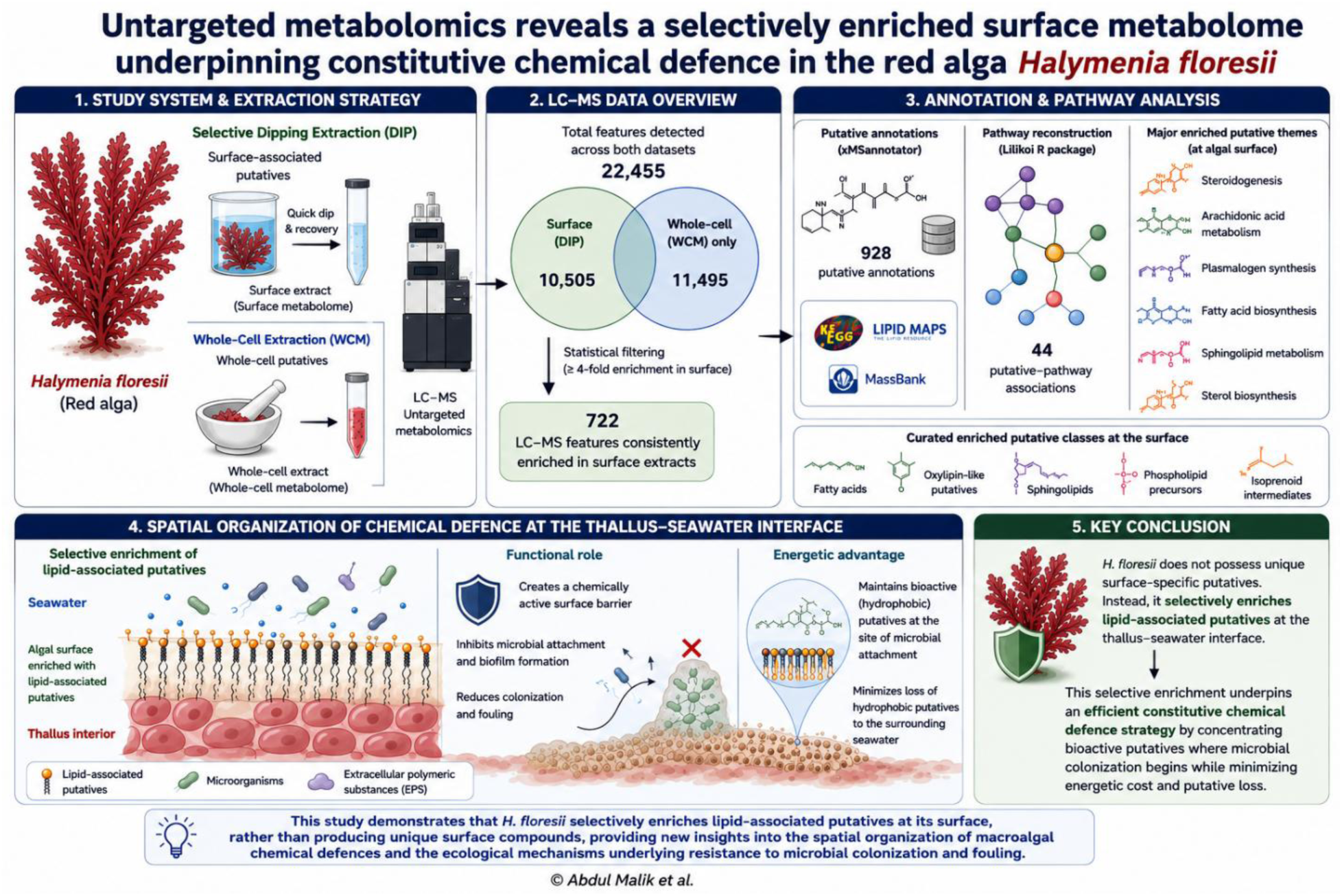

## 1. Introduction

Marine macroalgae thrive in highly dynamic coastal environments where their surfaces are continuously exposed to complex microbial communities, epiphytic algae, invertebrate larvae and other fouling organisms (Ren et al., 2022; Qian et al., 2022). The establishment of these organisms on algal surfaces can reduce light availability, impair nutrient exchange, increase drag, compromise reproductive performance and ultimately decrease host fitness (Qian et al., 2022). Consequently, marine macroalgae have evolved diverse physical and chemically mediated defence mechanisms that regulate interactions with surrounding organisms while maintaining beneficial microbial associations (Steinberg & de Nys, 2002; Egan et al., 2014; Abdul Malik et al., 2020a). Among these strategies, the production and localization of secondary metabolites at the thallus surface represent one of the most effective mechanisms for preventing microbial colonization and the establishment of both microfouling and macrofouling communities.

Unlike terrestrial plants, marine macroalgae function in an aqueous environment where soluble metabolites are rapidly diluted and dispersed into the surrounding seawater (Harlin & Rice, 1987; Steinberg & de Nys, 2002; Steinberg et al., 2002). Consequently, the effectiveness of chemical defence depends not only on metabolite production but also on maintaining sufficiently high concentrations of bioactive compounds at the thallus–seawater interface while minimizing continual metabolite loss and the metabolic costs of replenishment. Consistent with this strategy, several chemically defended macroalgae selectively localize defensive compounds at the thallus surface or within specialized storage structures, thereby maximizing protection precisely where interactions with microorganisms, epiphytes and herbivores are initiated (de Nys et al., 1998; Dworjanyn et al., 1999; Paul et al., 2006; Semmouri et al., 2024). Such spatial allocation is consistent with the Optimal Defence Theory (ODT), which predicts that defensive metabolites are preferentially allocated to tissues or regions experiencing the greatest probability of attack or contributing most strongly to organismal fitness, thereby maximizing defensive efficacy while minimizing metabolic investment (McKey, 1974; Rhoades, 1979; Cronin & Hay, 1996; Paul & Puglisi, 2004).

Marine red algae (Rhodophyta) are among the richest producers of structurally diverse natural products, including halogenated metabolites, terpenoids, sterols, fatty acids, oxylipins and other lipid-derived compounds that exhibit antimicrobial, antifouling, antiherbivore and antioxidant activities (Holdt & Kraan, 2011; Blunt et al., 2018). Chemically mediated defence has been extensively investigated in red macroalgae such as *Delisea pulchra*, *Laurencia* spp., *Asparagopsis* spp. and *Bonnemaisonia hamifera*. In these taxa, halogenated and other secondary metabolites stored in gland cells, released to the algal surface, or present at ecologically relevant concentrations can inhibit bacterial attachment and colonization, disrupt biofilm development, and deter herbivore feeding. (Maximilien et al., 1998; Dworjanyn et al., 1999; Vairappan et al., 2001; Paul et al., 2006 a,b; Nylund et al., 2008; Persson et al., 2011). Increasing evidence indicates that lipid-derived metabolites, particularly polyunsaturated fatty acids and oxylipins, function as signalling molecules involved in wound responses, microbial interactions and chemical defence in marine macroalgae (Bouarab et al., 2004; Ritter et al., 2008; Nylund et al., 2011; Barbosa et al., 2016;). Despite these advances, the spatial organization of surface-associated metabolites and their contribution to constitutive chemical defence remain poorly understood for most tropical red macroalgae.

Addressing this knowledge gap requires analytical approaches capable of comprehensively characterizing chemically complex and low-abundance surface-metabolites. Untargeted liquid chromatography–mass spectrometry (LC–MS) enables the simultaneous detection of thousands of metabolites without prior knowledge of their identities and has become one of the most sensitive approaches for characterizing complex natural-product mixtures (Fiehn, 2002; Patti et al., 2012). Combined with computational annotation platforms and pathway reconstruction, untargeted metabolomics allows metabolite classes and metabolic pathways to be inferred even when definitive structural identification is not yet possible. These approaches have been increasingly applied to marine macroalgae to investigate physiological responses to environmental stress, ecological interactions and secondary metabolite diversity (Kumar et al., 2016; Gaubert et al., 2019; Wahyuni et al., 2026). However, relatively few studies have specifically examined the surface metabolome, where ecological interactions between the host and its surrounding microbial community are initiated.

*Halymenia floresii* (Clemente) C.Agardh, 1817 (Rhodopyta, Halymeniales) is a tropical red macroalga widely distributed along the Yucatán Peninsula and is recognized as a valuable source of λ-carrageenan with considerable potential for aquaculture and human consumption. Despite its ecological and commercial importance, relatively little is known regarding its secondary metabolism or chemical ecology. Existing studies have largely focused on pigment composition under different environmental conditions (Godínez-Ortega et al., 2008) and the isolation of halymeniaol, a bioactive sterol derivative with antimalarial activity (Meesala et al., 2018). Interestingly, during experimental cultivation, *H. floresii* consistently exhibited remarkably clean thalli that remained largely free of bacterial biofilms, epiphytes and fouling organisms despite nutrient-rich culture conditions. These observations suggested the presence of an effective constitutive chemical defence associated with the algal surface (Abdul Malik et al. 2020a; 2020b).

In our previous study, surface-associated metabolites were selectively extracted using the non-destructive dipping (DIP) method developed by de Nys et al. (1998), allowing recovery of metabolites present at the thallus surface without disrupting underlying tissues. Those surface extracts inhibited quorum sensing in several Gram-negative marine bacteria naturally associated with *H. floresii*, including species of *Vibrio*, *Tenacibaculum*, *Maribacter* and *Aquimarina* (Abdul Malik et al., 2020b). Because quorum sensing regulates bacterial communication, biofilm development and the recruitment of fouling organisms, these findings suggested that *H. floresii* employs chemically mediated interference with bacterial signaling as an important antifouling strategy. Preliminary LC–MS fingerprinting further indicated the presence of several putative bioactive metabolites, including brominated compounds, volatile halogenated organic compounds, diterpenes and flavonoids (Abdul Malik et al., 2020b), although the broader chemical composition of the surface metabolome remained unresolved.

Although these findings strongly suggested the existence of an ecologically relevant surface metabolome, comprehensive chemical characterization remained challenging because only minute quantities of surface-associated metabolites could be recovered. Previous optimization of the non-destructive DIP extraction yielded approximately 600 ng cm⁻² of surface-associated metabolites from nearly 60 g of fresh algal biomass, substantially less than the corresponding whole-cell extracts. This limited quantity precluded conventional bioassay-guided purification and confirmatory structural identification by repeated MS/MS analysis and authentic standards. We therefore employed untargeted LC–MS metabolomics to identify molecular features preferentially localized to the algal surface and generate putative metabolite and pathway assignments without prior compound isolation. Complementary gas chromatography–mass spectrometry (GC–MS) analysis, with putative identification based on electron-impact spectral matching against the NIST library, provided an orthogonal approach for characterizing volatile and semi-volatile non-polar compounds. Together, these complementary analytical approaches were designed to overcome the limited quantity of surface extracts while providing the strongest feasible chemical evidence for metabolites potentially underlying the previously observed quorum-sensing inhibitory and antifouling activities of *H. floresii*.

The present study extends these preliminary observations by applying an untargeted LC–MS metabolomics approach to compare selectively extracted surface-associated metabolites (DIP) with corresponding whole-cell metabolite (WCM) extracts of *H. floresii*. Rather than searching exclusively for unique surface metabolites, we hypothesized that *H. floresii* maintains a selectively enriched surface metabolome that functions as an energetically efficient constitutive chemical defence, providing persistent protection at the thallus–seawater interface while minimizing metabolite loss and the energetic costs of defence. To test this hypothesis, untargeted LC-MS metabolomics was integrated with statistical filtering, putative metabolite annotation, pathway reconstruction and complementary GC–MS analysis to describe the metabolite classes and biochemical pathways preferentially enriched at the algal surface. By integrating these metabolomic data with our previous demonstration of quorum-sensing inhibition, this study provides the first pathway-level characterization of the surface metabolome of *H. floresii* and advances our understanding of the chemical ecology underpinning constitutive antifouling defence in marine macroalgae.

## 2. Materials and Methods

### 2.1. Algal Material and extractions

*H. floresii* were cultivated in Centro de Investigación y de Estudios Avanzados del Instituto Politécnico Nacional (Cinvestav), Yucatán, México under controlled environmental conditions in acrylic cultivation cylinders inside a cultivation chamber after several weeks of acclimatization. The cultures used natural seawater, filtered (0.5 micron) and UV-sterilized (UV-filter 0.4 micron) at 30 psu, 22 ± 2° C and pH 8.0 ± 0.3 under continuous aeration at a 12/12 light/dark cycle with an intensity of 110 ± 7.2 µmol photons m^-2^ s^-1^. Every week *H. floresii* was pulse-fed with Provasoli Enriched Seawater Media (0.3%) for 24 h. After eight weeks of cultivation, approximately 60 g fw were harvested and extracted (∼ 5 g for each time period in duplicates). In order to selectively extract the surface-associated metabolites, samples were taken for the DIP extraction as previously reported (A Abdul Malik et al., 2020b) since the use of n-hexane allows to extract the surface metabolites of *H. floresii* without lysis of the cortical cell wall and with a satisfying reproducibility. Thus, the surface metabolites were gently and selectively extracted using n-hexane, a non-polar solvent, using the dipping method (DIP) previously described by de Nys et al. (1998).

Briefly, *H. floresii* thalli duplicates were dipped into n-hexane for six durations of 10, 20, 30, 40, 50 and 60 seconds under stirring, and the resulting extracts were pooled together. After the DIP extraction, *H. floresii* thalli were rinsed with sterile seawater (30 psu) and individually freeze-dried. The lyophilized samples were then exhaustively extracted with n-hexane by maceration for 24 h under agitation at room temperature to yield the Whole Cell Metabolites fraction (WCM). For comparison purposes, the same solvent was used. The extracts were then filtered through Whatman filter to remove any particles, and the solvent was evaporated under vacuum. Henceforth, the surface metabolites and whole-cell metabolites will be referred to as DIP and WCM, respectively.

### 2.2. LC-MS conditions and data pre-processing

DIP and WCM extracts were analyzed by LC-MS as described in Abdul Malik et al. (2020b). Briefly, the chemical profiling of *H. floresii* extracts were analyzed by LC/MS using an LC-ESI-Q-TOF-MS (Dionex, Ultimate 3000, Bruker, micrOTOF-QII) system (Bruker Daltonik GmbH, Bremen, Germany). Separations were performed using an analytical reversed-phase column. A sample of 10 µL (in triplicates) of the extract was injected for every analysis at a flow rate of 0.5 mL min^-1^ with a column temperature of 30 °C and the elution gradient was adopted from Othmani et al. (2016). Binary programming of methanol (MeOH) and water (H_2_O) was injected at a linear gradient from MeOH/H_2_O (40:60, v/v) to MeOH/H_2_O (85:15, v/v) in 3 min and to 100% MeOH in 4 min, followed by a final isocratic step with 100% MeOH for 18 min, finishing by a return to the initial conditions (0.1 min) and equilibration of the column (9.9 min). Regarding the mass spectrometer, operating conditions were set as following: drying temperature: 350 °C, capillary voltage: 4 kV, nebulizer pressure: 3.45 bars, drying gas: helium at a flow rate of 12 L min^-1^ (Othmani et al., 2016). Mass spectra acquisition was set to 0.5 Hz from m/z 50 to 1,000.

The raw data files exported as “.mzXML” format from the instrument were used for the analyses. The LC/MS raw data in.mzXML format were uploaded into the XCMS Online platform (https://xcmsonline.scripps.edu/). The DIP LC/MS data in duplicates were uploaded to the assigning a ‘Single’ job treatment whereas the WCM LC/MS data were either assigned a ‘Multi’ job as the WCM data included six extraction periods of 10, 20, 30, 40, 50 and 60 seconds in duplicates or a ‘Single’ job which included individual sets. Both data sets were treated with the same default parameters, UPLC / Bruker Q-TOF pos, with mzdiff of ‘0.01’ and SNR (signal-noise ratio) threshold of ‘6’. The method used for the feature detection is ‘centWave’ with 10 ppm of m/z tolerance and minimum/maximum peak width of 5 seconds/20 seconds. The *p*-value threshold was set between 0.001 (highly significant) to 0.05 (not significant). After several steps of treatment such as filtration, retention time correction, fill-in missing peak, and annotation of adducts and isotopes, the pre-processed data were downloaded from the platform.

### 2.3 Statistical Analysis

Statistical analyses were performed in R (R Core Team, Vienna, Austria, 2025), an open-source environment widely used for statistical computing, data visualization and metabolomic data analysis. The LC–MS feature matrix, comprising m/z, retention time (RT) and peak intensity values for all biological replicates, was imported into R for downstream processing and comparative analysis (Smith et al., 2006). The dataset (HF.xlsx; Supplementary Table S1) included two surface-associated extract samples (DIP: Hf1 and Hf2) and six whole-cell extract samples (WCM: Hf3–Hf8).

The analytical workflow comprised nine sequential steps: Step 1: Importing the dataset (here, HF.xlsx, available in Supplementary section Table S1) which contains the mzmax, rtmax and intensities (surface metabolites: Hf1, Hf2; whole-cell metabolites: Hf3, Hf4, Hf5, Hf6, Hf7 and Hf8). Step 2: Question Zero: The peaks either present or absent in the DIP and WCM datasets were managed in this step by indicating NA (Not Available) in the respective columns (as they were filled by ‘0’) (detailed in the Results section and also see, HF.xlsx dataset in the Supplementary section Table S1). Step 3: Defining experimental subsets: Experimental subsets were defined here with mz, rt and peaks intensity as their variables. Variables mz and rt were merged and then the DIP, WCM subsets were defined and aligned into separate columns. Step 4: Remove the duplicate peaks: Duplicate peaks were discarded from the defined subsets, and the unique peaks were kept for further analysis. Step 5: Merging the subsets into one cluster group: DIP subsets (Hf1 and Hf2) and WCM subsets (Hf3, Hf4, Hf5, Hf6, Hf7 and Hf8) were merged into respective cluster groups. Step 6: Reduction of noise/incertitude: Only the peaks present in all the subsets were selected which means the NA peaks were discarded, and the incertitude will be reduced drastically. Step 7: m/z ‘common’ in DIP and WCM: Only the peaks which were common in DIP and WCM were selected and merged as ‘x’ and ‘y’ variables. Step 8: Log2fold change in intensities: Among the common peaks selected previously, log2fold change was calculated in order to scrutinize the results tightly. Step 9: Plotting the results: The results were plotted graphically using R package.

Missing-value handling and complete-case feature filtering were undertaken with consideration of established approaches for untargeted metabolomics data (Wei et al., 2018), while relative differences in feature abundance were evaluated using log₂ fold-change analysis, as commonly applied in LC–MS-based metabolomics. The R scripts used for the analysis will be made available through a public GitHub repository upon publication.

### 2.4. Metabolite Annotation and Pathway Mapping

After statistical analysis, the list of common features, consisting of accurate mass-to-charge ratio (m/z) and retention time (RT) values were used as input for annotation. Prior to analysis, duplicate features were removed and only unique m/z–RT pairs were retained. Putative metabolite annotations were performed using the R package xMSannotator, which includes utilities for querying online databases such as ChemSpider, KEGG, HMDB, and LipidMap (Uppal et al., 2017) in positive ionization mode. Accurate mass matching was performed with a mass tolerance of 10 ppm, and common positive-ion adducts ([M+H]+, [M+Na]+ and [M+NH4]+) were considered during the annotation process. Database searches were executed using four computational threads. For each detected feature, xMSannotator generated candidate metabolites together with their corresponding HMDB identifiers, molecular formulas, monoisotopic masses and adduct assignments.

In order to facilitate the biological understanding of the annotations and to assess the relevance of putative metabolites to marine macroalgae, the putative candidates were subsequently cross-referenced against complementary metabolomic resources including the KEGG (Kanehisa et al., 2023), LIPID MAPS (Fahy et al., 2009), MassBank (Horai et al., 2010). These databases were employed to validate the molecular formulae, chemical classification, lipid assignments and the presence of known metabolites and pathway associations whenever available. Such cross-referencing provided further evidence for metabolite class assignment and facilitated the recognition of compounds previously reported for marine organisms and algae. Further, additional metabolite descriptors, including structural and biological information available were retrieved and associated with each putative annotation. Subsequently, metabolites were mapped to biochemical pathways employing Lilikoi package (Huang et al., 2018) pathway enrichment process. Metabolites–pathway associations were extracted to identify the most represented metabolic pathways in the metabolome of *H. floresii*.

Since the metabolite annotations were mainly based on accurate-mass matching and database search without any confirmation using authentic standards or tandem mass spectrometry (MS/MS), all the identities were considered ‘putative annotations’ and belong to Metabolomics Standards Initiative (MSI) Level 4 confidence (Sumner et al., 2007). Therefore, annotations should be interpreted as candidate metabolites and metabolite classes rather than definitive compound identifications.

### 2.3 GC-MS conditions

The DIP and WCM extracts were analysed by gas chromatography (Agilent, Model 7890B, Palo Atlo, CA, USA) coupled with mass spectrometry (Agilent, Model 7000C, Palo Alto, CA, USA). A sample of 2 µL of each DIP extract obtained was injected at temperature of 250° C under mode splitless. Helium as used as a carrier gas at a flow rate of 1mL/min. The column temperature was programmed as follows: initial temperature at 100° C for 4 min, ramps of 10° C/min to 300° C for 15 min. Mass detector conditions were electron impact mode at 70 eV, transfer line temperature 270° C, source temperature 230° C, mass acquisition range 50 – 500 amu (atomic mass units) and solvent delay 3.75 min. The compounds were identified by comparison of their mass spectra with those from the NIST/EPA/NIH Mass Spectral Library version NIST 2.2 (Agilent, Palo Alto, CA, USA).

## 3. Results

### 3.1. Comparative metabolomic fingerprint of surface-associated and whole-cell metabolites

Untargeted LC-MS analysis generated comprehensive metabolomic fingerprints for both the selectively extracted surface-associated metabolites (DIP) and the corresponding whole-cell metabolite (WCM) fractions of *H. floresii*. Following feature detection and preprocessing in XCMS Online, a total of 22,455 LC-MS features (see Supplementary Section, Table S1) were detected across all samples. Among these, 10,505 m/z features (see Supplementary Section, Table S2) were detected in the surface-associated (DIP) extracts, whereas the whole-cell metabolite (WCM) extracts contained an additional 11,495 features that were not detected in the surface extracts. Thus, the whole-cell metabolome represented a broader chemical inventory, while the surface metabolome comprised a subset of metabolites exposed at the algal surface.

Comparison of the DIP and WCM datasets revealed marked differences in metabolite abundance and enrichment where numerous high-intensity LC-MS features displayed pronounced enrichment in the DIP surface extracts. As illustrated by the scatter plot of peak intensity against log₂ fold change (Fig. 1), features positioned at approximately 0e+00 on the log₂ fold-change axis exhibited comparable peak intensities in DIP and WCM extracts, indicating little or no relative enrichment in either fraction. In contrast, features with positive log₂ fold-change values were more abundant in DIP extracts, whereas those with negative values were relatively more abundant in WCM extracts. The distribution of features across these regions demonstrates that the surface-associated metabolome is characterized not simply by the presence of particular compounds but by differences in their relative abundance between the two fractions. These observations indicate that *H. floresii* selectively concentrates specific metabolites at its surface while maintaining a much larger metabolite reservoir within the algal tissues.

**Fig. 1.**
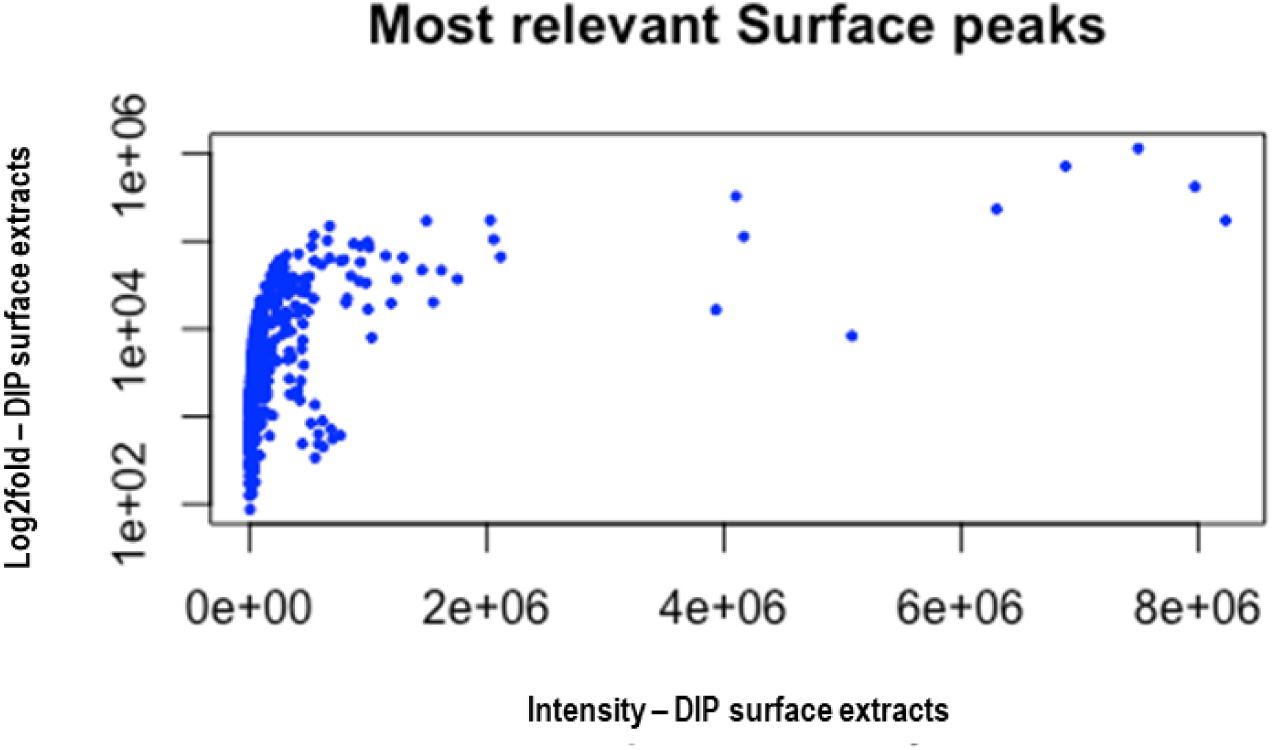
Most relevant surface-associated metabolites identified after log2 fold-change analysis. Peak intensities in the DIP fraction are plotted against enrichment values. Peaks at 0e+00 correspond to metabolites with similar intensities in the DIP and WCM fractions.

### 3.2 Statistical identification of surface-enriched metabolites

To identify metabolites involved in surface-associated defence, LC-MS features were subjected to a stringent statistical filtering workflow implemented in R. Duplicate m/z – retention time pairs were removed, and only features consistently detected throughout all extraction periods were retained. This conservative approach minimized analytical variability and ensured that subsequent comparisons were based exclusively on reproducible features. Relative metabolite abundance was evaluated calculating the log2 fold-change between DIP and WCM intensities. The distribution of log2 fold-change values approximated a normal distribution centered near zero (Fig. 2), indicating the most metabolites occurred at comparable abundances in both extracts. However, a threshold of log2(DIP/WCM) >2 corresponded to the features occurring at least four-fold higher concentration in the surface extracts than in the whole-cell extracts.

**Fig. 2.**
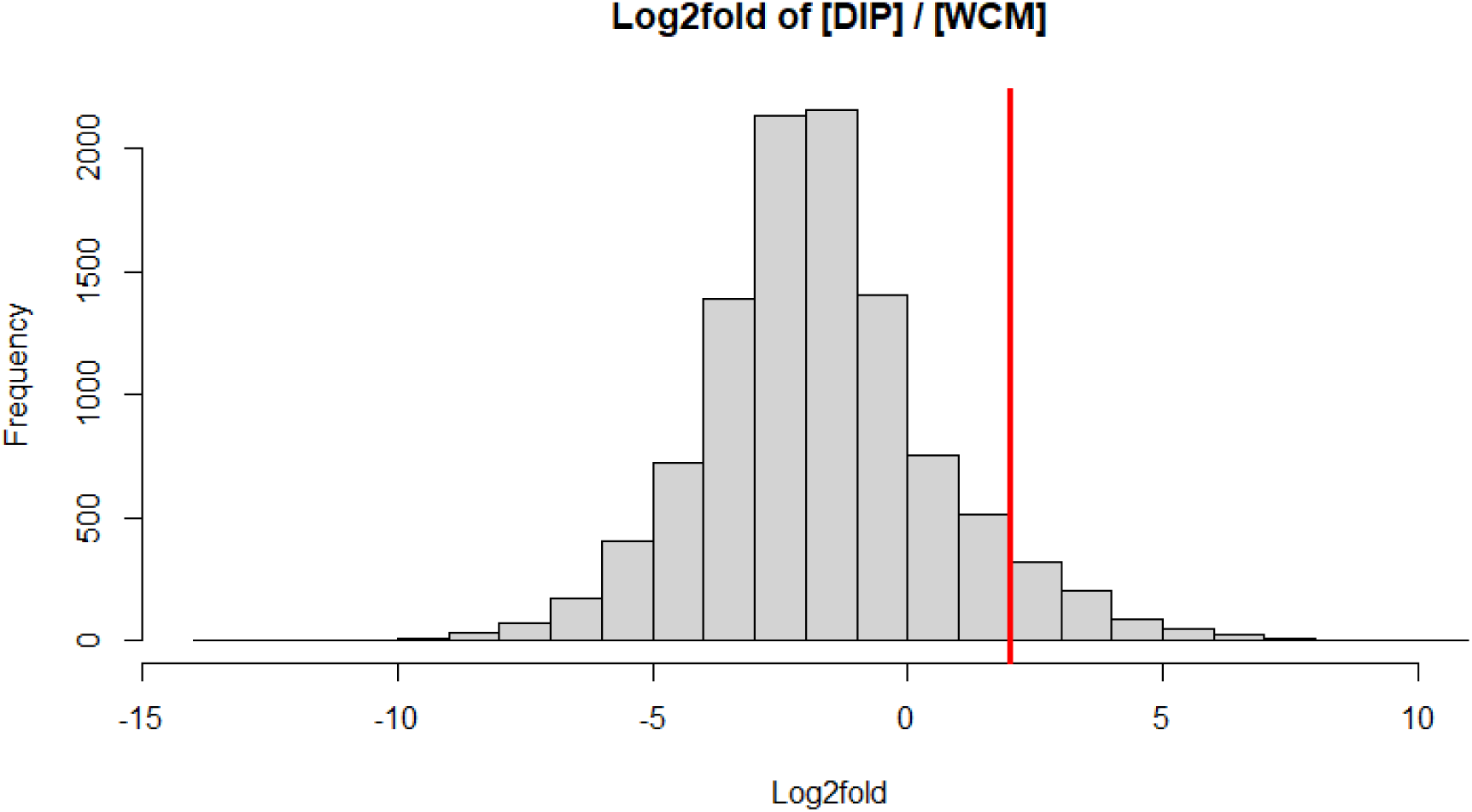
Distribution of log2(DIP/WCM) values for all common LC–MS features. The red vertical line indicates the selection threshold (log2 fold-change > 2), corresponding to metabolites at least four-fold more abundant in the surface-associated fraction (DIP).

Following removal of peaks not consistently detected throughout the extraction periods, 722 LC-MS features (see Supplementary Section Table S3) remained that exhibited at a four-fold higher abundance in the DIP extracts relative to the WCM extracts (Fig. 3). These features were distributed across the entire detected mass range (approximately m/z 70 - 500).

**Fig. 3.**
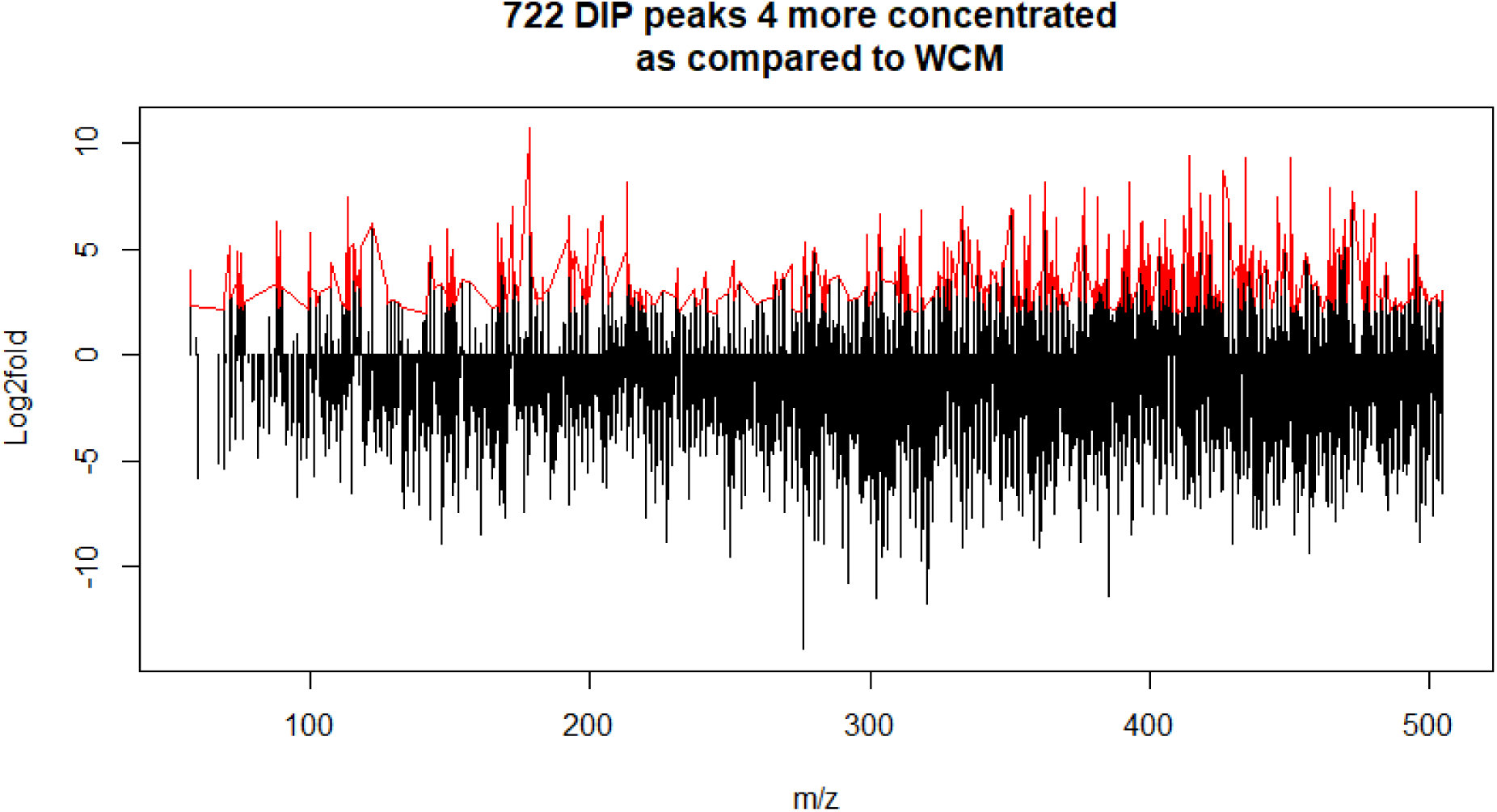
Distribution of log2 fold-change values across the detected m/z range. Red peaks indicate metabolites enriched at the algal surface-associated fraction. A total of 722 features exceeded the four-fold enrichment threshold.

### 3.3. Putative metabolite annotation

The 722 surface-enriched LC-MS features were subsequently subjected to metabolite annotation using xMSannotator. Considering multiple ions adducts within a mass tolerance of 10 ppm, the annotation workflow generated 928 putative metabolite annotations (see Supplementary Section, Table S4), reflecting multiple candidate assignments for individual LC-MS features. Cross-referencing against KEGG, LIPID MAPS and MassBank enabled assignment of metabolite classes and facilitated biological interpretation. Metabolite identities remain putative (MSI Level 4) focused primarily on metabolite classes and metabolic pathways rather than individual compound identities. Among the annotated metabolites, a curated subset comprised chemically plausible macroalgal metabolites including saturated and polyunsaturated fatty acids, oxylipin-like compounds, sphingolipids, phospholipid precursors, amino acid derivatives and isoprenoid intermediates (Table 1). Conversely, several annotations were regarded as annotation artefacts generated by database matching and were excluded from the biological interpretation.

**Table 1.** Differential metabolites putatively identified in *Halymenia floresii* using untargeted liquid chromatography–mass spectrometry (LC–MS)-based metabolomics.

| S.No | Class of chemical | Observed ion (m/z) | Ion form | t<sub>R</sub> (min) | Formula | Theoretical ion (m/z) | ppm | log2Fold | Confidence level | Putative identity |
| --- | --- | --- | --- | --- | --- | --- | --- | --- | --- | --- |
| 1 | Polyunsaturated fatty acid | 333.2787 | (M+H)+ | 21.30 | C22H36O2 | 333.2788 | -3.3653 | 6.96 | 2 | Adrenic acid |
| 2 | Very long-chain PUFA | 374.3060 | (M+NH4) + | 12.92 | C24H36O2 | 374.3054 | 1.7102 | 2.87 | 2 | Tetracosahexaenoic acid |
| 3 | Saturated fatty acid | 302.3061 | (M+NH4) + | 28.31 | C18H36O2 | 302.3054 | 2.3212 | 3.05 | 2 | Stearic acid |
| 4 | Fatty acid | 199.1684 | (M+H)+ | 12.86 | C12H22O2 | 199.1693 | -4.1833 | 5.92 | 2 | trans-Dodec-2-enoic acid |
| 5 | Hydroxy fatty acid | 133.0865 | (M+H)+ | 13.66 | C6H12O3 | 133.0859 | 4.2395 | 2.20 | 2 | (R)-3-Hydroxyhexanoic acid |
| 6 | Oxylipin-like metabolites | 352.2514 | (M+NH4) + | 12.30 | C20H30O4 | 352.2482 | 8.9983 | 2.04 | 3 | Prostaglandin A2/J2/B2-like metabolites |
| 7 | Oxylipin-like metabolites | 384.2408 | (M+NH4) + | 15.47 | C20H30O6 | 384.2381 | 7.1324 | 2.24 | 3 | Leukotriene-like metabolites |
| 8 | Amino acid derivative | 251.0612 | (M+H)+ | 13.16 | C8H14N2O5S | 251.0696 | -33.458 | 4.43 | 2 | γ-Glutamylcysteine |
| 9 | Phospholipid precursor | 464.0923 | (M+NH4) + | 16.57 | C11H20N4O11P2 | 464.0942 | -4.0482 | 2.59 | 2 | CDP-Ethanolamine |
| 10 | Fatty alcohol | 293.2813 | (M+Na)+ | 20.51 | C18H38O | 293.2815 | -0.5556 | 2.55 | 2 | Octadecanol |
| 11 | Sphingolipid | 302.3061 | (M+H)+ | 28.31 | C18H39NO2 | 302.3054 | 2.3143 | 3.05 | 2 | Sphinganine |
| 12 | Isoprenoid precursor | 268.9963 | (M+NH4) + | 12.80 | C5H12O7P2 | 268.9950 | 4.8021 | 3.78 | 2 | Dimethylallyl pyrophosphate (DMAPP) |
| 13 | Isoprenoid precursor | 268.9963 | (M+NH4) + | 12.80 | C5H12O7P2 | 268.9950 | 4.8021 | 3.78 | 2 | Isopentenyl pyrophosphate (IPP) |
| 14 | Isoprenoid-like compounds | 304.2660 | (M+Na)+ | 13.75 | C20H30O | 304.2635 | 8.1407 | 3.59 | 3 | Vitamin A / Retinol-like metabolites |
| 15 | Sulfonic amino acid | 143.0485 | (M+NH4) + | 14.03 | C2H7NO3S | 143.0485 | -0.1553 | 5.15 | 2 | Taurine |
| 16 | Acylcarnitine | 400.3400 | (M+H)+ | 13.65 | C23H45NO4 | 400.3421 | -5.2578 | 4.74 | 2 | L-Palmitoylcarnitine |

Metabolites are grouped according to their chemical class and are presented with their observed mass-to-charge ratio (m/z), ionization adduct, retention time (t<sub>R</sub>, min), molecular formula, theoretical m/z, mass error (ppm), log2 fold change, metabolite identification confidence level, and putative identity. Metabolite annotation was performed using accurate mass measurements and spectral matching against available metabolomics databases. Identification confidence was assigned according to the Metabolomics Standards Initiative (MSI) guidelines, where Level 2 represents putatively annotated compounds based on accurate mass similarity to reference databases without confirmation using authentic standards, and Level 3 represents putatively characterized compound classes based on diagnostic physicochemical properties and spectral features. Positive log2 fold-change values indicate metabolites with higher relative abundance in the experimental group compared with the reference group.

**Table 2.** Metabolic pathways assigned to surface-enriched putative metabolites identified in *H. floresii* and their proposed ecological relevance.

| S.No | Putative identity | Assigned pathway | Proposed biological role | Evidence in macroalgae | Key literature |
| --- | --- | --- | --- | --- | --- |
| 1 | Adrenic acid | α-Linolenic/linoleic acid metabolism | Membrane lipid; precursor of signalling oxylipins | Yes | Godínez-Ortega et al. (2012); Carpena et al. (2024) |
| 2 | Tetracosahexaenoic acid | α-Linolenic/linoleic acid metabolism | Membrane fluidity and lipid metabolism | Likely | Carpena et al. (2024) |
| 3 | Stearic acid | Plasmalogen synthesis | Structural membrane lipid | Yes | Godínez-Ortega et al. (2012); Carpena et al. (2024) |
| 4 | trans-Dodec-2-enoic acid | Fatty acid biosynthesis | Antimicrobial fatty acid | Likely | Goecke et al. (2010); Carpena et al. (2024) |
| 5 | (R)-3-Hydroxyhexanoic acid | Fatty acid biosynthesis | Lipid turnover | Likely | Carpena et al. (2024) |
| 6 | Prostaglandin A2/J2/B2-like metabolites | Arachidonic acid metabolism | Defence signalling, wound response, antimicrobial activity | Oxylipin class reported | Nylund et al. (2011); Goulitquer et al. (2012) |
| 7 | Leukotriene-like metabolites | Arachidonic acid metabolism | Stress signalling | Oxylipin class reported | Nylund et al. (2011); Goulitquer et al. (2012) |
| 8 | γ-Glutamylcysteine | Cysteine metabolism | Glutathione biosynthesis; antioxidant defence | Likely | Kumar et al. (2016); Carpena et al. (2024) |
| 9 | CDP-Ethanolamine | Plasmalogen synthesis | Membrane phospholipid biosynthesis | Likely | Kumar et al. (2016); Carpena et al. (2024) |
| 10 | Octadecanol | Plasmalogen synthesis | Lipid metabolism | Likely | Carpena et al. (2024) |
| 11 | Sphinganine | Sphingolipid metabolism | Membrane signalling and stress adaptation | Yes | Kumar et al. (2016); Carpena et al. (2024) |
| 12 | Dimethylallyl pyrophosphate (DMAPP) | Steroid biosynthesis | Terpenoid and sterol biosynthesis | Universal metabolite | Meesala et al. (2018); Carpena et al. (2024) |
| 13 | Isopentenyl pyrophosphate (IPP) | Steroid biosynthesis | Biosynthesis of terpenoids and sterols | Universal metabolite | Meesala et al. (2018); Carpena et al. (2024) |
| 14 | Vitamin A / Retinol-like metabolites | Retinol metabolism | Antioxidant metabolism | Carotenoid pathway analogue | Godínez-Ortega et al. (2012); Carpena et al. (2024) |
| 15 | Taurine | Taurine and hypotaurine metabolism | Osmoregulation; antioxidant defence | Yes | Kumar et al. (2016); Carpena et al. (2024) |
| 16 | L-Palmitoylcarnitine | Fatty acid metabolism | Fatty acid transport | Possible | Kumar et al. (2016) |

The table summarizes the putative metabolite identity, the corresponding metabolic pathway assigned by Lilikoi, the proposed biological role, evidence supporting its occurrence in macroalgae (Yes = experimentally reported; Likely = chemically or biosynthetically plausible based on related taxa or metabolite class; Possible = inferred from conserved metabolic pathways but lacking direct evidence), and representative literature supporting each assignment. Because metabolite identities were inferred from accurate-mass matching and pathway reconstruction, all compound and pathway assignments should be regarded as putative and interpreted at the metabolite-class and pathway levels rather than as definitive metabolite identifications.

### 3.4 Lipid associated metabolic pathways dominate the surface-enriched metabolome

Pathway reconstruction using Lilikoi identified 44-metabolite-pathway associations (see Supplementary section, Table S5) derived through the xMSannotator program. Although pathway mapping putative annotations, the majority of biologically plausible metabolites converged on lipid-associated pathways. The most represented pathways included steroidogenesis (8 matches), arachidonic acid metabolism (6 matches), bile acid biosynthesis (5 matches), plasmalogen synthesis (3 matches), retinol metabolism (3 matches), α-linolenic acid and linoleic acid metabolism (2 matches), fatty acid biosynthesis (2 matches), and steroid biosynthesis (2 matches). Additional pathways represented by single metabolites included the citric acid cycle, cysteine metabolism, galactose metabolism, gluconeogenesis, pyrimidine metabolism, sphingolipid metabolism, taurine and hypotaurine metabolism, and fatty acid metabolism (see Supplementary Section, Table S5).

Surface-enriched putative metabolites identified by untargeted LC–MS metabolomics were assigned to metabolic pathways using the Lilikoi R package, which maps annotated metabolites to KEGG pathways based on putative metabolite identities. Following pathway assignment, each metabolite was manually curated through cross-referencing with the scientific literature to evaluate its reported occurrence, biochemical function and ecological relevance in marine macroalgae.

Mass spectra were interpreted by comparing observed m/z values with the theoretical masses of the expected ions and the curated putative dataset demonstrated that the most biologically plausible annotations corresponded predominantly to lipid-derived metabolites, including polyunsaturated fatty acids (m/z 333.2787, m/z 374.3060), oxylipin-like compounds (m/z 352.2514, m/z 384.2408), sphingolipids (m/z 302.3061), phospholipid precursors (m/z 464.0923) and isoprenoid intermediates (m/z 268.9963) (Table 1). Representative putatives included adrenic acid (m/z 333.2787, RT 21.30), tetracosahexaenoic acid (m/z 374.3060, RT 12.92), stearic acid (m/z 302.3061 RT 28.31), trans-Dodec-2-enoic acid (m/z 199.1684, RT 12.86), (R)-3-Hydroxyhexanoic acid (m/z 133.0865, RT 13.66), Prostaglandin A2/J2/B2-like metabolites (m/z 352.2514, RT 12.30), leukotriene-like metabolites (m/z 384.2408, RT 15.47), γ-Glutamylcysteine (m/z 251.0612, RT 13.16), CDP-ethanolamine (m/z 464.0923, RT 16.57), octadecanol (m/z 293.2813, RT 20.51), sphinganine (m/z 302.3061, RT 28.31), dimethylallyl pyrophosphate (m/z 268.9963, RT 12.80), isopentenyl pyrophosphate (m/z 268.9963, RT 12.80), taurine (m/z 143.0485, RT 14.03), and L-palmitoylcarnitine (m/z 400.3400, RT 13.65).

### 3.5. GC–MS analysis independently confirms the predominance of lipid-derived metabolites

To complement the untargeted LC–MS metabolomic analysis, selected extracts were analysed by GC–MS to characterize volatile and semi-volatile metabolites amenable to gas chromatographic separation. Compound identification was performed by comparison of electron-impact mass spectra with the NIST/EPA/NIH Mass Spectral Library (NIST 2.2). GC–MS results putatively annotated for twenty-two compounds belonging predominantly to long-chain fatty acid esters, unsaturated fatty acid derivatives, terpenoid alcohols and aliphatic hydrocarbons (Table 3). The most frequently detected metabolites included hexadecanoic acid ethyl ester, methyl palmitate, methyl stearate, methyl oleate, methyl elaidate, methyl arachidonate, arachidonic acid ethyl ester and the methyl ester of eicosapentaenoic acid. In addition, phytol, neophytadiene, phytane and tetramethylhexadecenol were identified, together with several long-chain hydrocarbons including n-eicosane and tetramethylheptadecane.

**Table 3.** Compounds putatively annotated in the surface extracts of *Halymenia floresii* by complementary GC–MS analysis and their proposed ecological relevance.

| S.No | Compounds | Parent class | Solvent | m/z | RT (min) | Ecological relevance |
| --- | --- | --- | --- | --- | --- | --- |
| 1 | Hexadecanoic acid, ethyl ester | Palmitic acid ester; saturated fatty-acid derivative | DCM | 88 | 16.2 | Membrane lipid, antimicrobial |
| 2 | Ethyl Oleate | Oleic acid ester; monounsaturated fatty-acid derivative | DCM | 96 | 17.8 | Antimicrobial, membrane integrity |
| 3 | Stearic acid, ethyl ester | Stearic acid ester; saturated fatty-acid derivative | DCM | 88 | 18 | Structural lipid |
| 4 | Methyl myristate | Myristic acid ester; saturated fatty-acid derivative | MeOH | 74.1 | 13.4 | Membrane lipid, antimicrobial defence |
| 5 | Methyl elaidate | Trans-C18:1 fatty-acid ester | MeOH | 55.1 | 17.2 | Hydrophobic surface constituent, membrane lipid |
| 6 | Methyl oleate | Oleic acid ester | MeOH | 55.1 | 17.3 | Antimicrobial fatty acid derivative; biofilm inhibition |
| 7 | 3,7,11,15-Tetramethyl-2-hexadecen-1-ol | Phytol | MeOH | 81 | 15.1 | Chlorophyll derived diterpenoid involved in stress tolerance |
| 8 | Methyl arachidonate | Arachidonic acid ester; C20:4 PUFA derivative | MeOH | 79 | 18.7 | Oxylipin precursor; defence signalling; antimicrobial |
| 9 | Arachidonic acid, ethyl ester | Arachidonic acid ester | DCM:MeOH | 79 | 18.7 | Oxylipin precursor |
| 10 | Phytol | Diterpenoid alcohol | DCM:MeOH | 81 | 15.1 | Antioxidant, antimicrobial |
| 11 | Methyl palmitoleate | Palmitoleic acid ester; C16:1 derivative | DCM:MeOH | 55.1 | 15.3 | Antimicrobial lipid; biofilm inhibition |
| 12 | Methyl 8-octadecenoate | C18:1 fatty-acid ester | DCM:MeOH | 55.1 | 17.2 | Hydrophobic surface constituent; antimicrobial |
| 13 | Isopropyl palmitate | Palmitic acid ester | DCM;<br>DCM:MeOH | 255.9;<br>101.9 | 16.5 | Enhances persistence of lipid barrier; reduces diffusional loss |
| 14 | Elaidic acid, methyl ester | Same as methyl elaidate | DCM;<br>DCM:MeOH | 55.1 | 17.2 | Structural lipid; hydrophobic barrier |
| 15 | Methyl stearate | Stearic acid ester | MeOH;<br>DCM:MeOH | 74.1 | 17.4 | Membrane stability; hydrophobic barrier |
| 16 | Methyl palmitate | Palmitic acid ester | MeOH;<br>DCM:MeOH | 74.1 | 15.5 | Defence against microbial colonization; Antimicrobial |
| 17 | Neophytadiene | Diterpene hydrocarbon | MeOH;<br>DCM:MeOH | 95 | 14.7 | Antimicrobial |
| 18 | 5,8,11,14,17-Eicosapentaenoic acid, methyl ester, (all-Z)- | EPA methyl ester; C20:5 PUFA derivative | MeOH;<br>DCM:MeOH | 79 | 18.7 | Oxylipin precursor |
| 19 | Phytane | Branched isoprenoid hydrocarbon | n-hexane | 71.1 | 16.1 | Hydrophobic surface constituent |
| 20 | n-Eicosane | Long-chain hydrocarbon | n-hexane | 71.1 | 16.2 | Hydrophobic surface barrier |
| 21 | Heptadecane, 2,6,10,15-tetramethyl- | Branched isoprenoid-like hydrocarbon | n-hexane | 85.1 | 16.4 | Protection against environmental stress |
| 22 | Ethyl elaidate | Trans-C18:1 fatty-acid ester | n-hexane | 55.1 | 17.8 | Antimicrobial activity |

Compounds detected in the surface-associated extracts of *Halymenia floresii* by gas chromatography–mass spectrometry (GC–MS) following extraction with dichloromethane (DCM), methanol (MeOH), DCM:MeOH (1:1, v/v) or n-hexane.
Compound identities were assigned by comparison of electron ionization (EI) mass spectra with the NIST Mass Spectral Library and are reported together with their parent chemical class, extraction solvent, characteristic diagnostic ion ( $m/z$ ), retention time (RT), and proposed ecological relevance.
Ecological functions were assigned through manual curation of the marine chemical ecology literature and primarily reflect roles reported for the corresponding metabolite classes rather than definitive biological activities of the individual compounds.

Most identified metabolites belonged to lipid-derived chemical classes, particularly saturated and unsaturated fatty acids, fatty acid esters and isoprenoid-derived compounds. Fatty acid derivatives were detected in both methanolic and dichloromethane extracts, whereas the non-polar n-hexane extracts contained predominantly hydrophobic hydrocarbons and long-chain lipid derivatives. This solvent-dependent distribution is consistent with the expected physicochemical properties of these metabolite classes and supports the selective recovery of lipophilic compounds by non-polar extraction solvents.

Importantly, the GC–MS results closely paralleled the untargeted LC–MS metabolomic analysis. Several compounds detected by GC–MS, including palmitic acid, stearic acid, arachidonic acid derivatives, phytol and isoprenoid metabolites, correspond to metabolite classes represented among the putative LC–MS annotations and mapped predominantly to fatty acid metabolism, arachidonic acid metabolism, steroidogenesis and other lipid-associated pathways. Although the analytical platforms target different subsets of the metabolome, both datasets independently indicate that lipid-derived metabolites constitute the dominant chemical component of the *H. floresii* surface metabolome.

## 4. Discussion

Metabolomics, a rapidly evolving discipline, applies advanced chromatographic, spectrometric and spectroscopic methods to characterize the collection of low-molecular-weight compounds associated with cellular metabolic pathways (Fernie et al., 2004; Patti et al., 2012; Carpena et al., 2024). Because metabolites constitute downstream products of gene regulation, enzymatic activity and environmental influences, metabolomic profiles can bridge the gap between changes in metabolic pathways and their associated phenotypic responses (Fiehn, 2002; Patti et al., 2012; Johnson et al., 2016). The present study extends our understanding of the chemical ecology of *H. floresii* by providing the first comprehensive characterization of its surface-associated metabolome. The combination of untargeted LC–MS metabolomics and complementary GC–MS analysis enabled the characterization of an otherwise analytically challenging, low-abundance surface extract. Collectively, the results reveal that the surface chemistry of *H. floresii* is dominated not by unique metabolites, but by the selective enrichment of lipid-associated metabolites, which are likely to play an important role in the species’ constitutive chemical defence (Park et al., 2023).

### 4.1 Surface enrichment represents a chemical ecology strategy

One of the principal findings of this study is that the distinction between the surface-associated (DIP) and whole-cell (WCM) metabolomes of *H. floresii* is driven primarily by metabolite abundance rather than metabolite identity. Although the whole-cell metabolome comprised a substantially larger chemical inventory, only a subset of metabolites was significantly enriched at the thallus surface, with abundances at least four-fold higher than in the corresponding whole-cell extracts. These findings indicate that *H. floresii* does not rely on a unique suite of surface-specific metabolites; instead, it selectively concentrates particular metabolites at the organism–environment interface. This spatial organization of metabolites is consistent with current concepts in marine chemical ecology, which recognize that macroalgae strategically localize defensive chemistry to sites of greatest ecological relevance (de Nys et al., 1998; Paul et al., 2006; Abdu Malik et al., 2020b). As the thallus surface is the primary interface for microbial colonization, epiphyte settlement, and herbivore interactions, localized metabolite enrichment maximizes defensive efficacy while minimizing the energetic costs of maintaining high concentrations of bioactive compounds throughout the thallus (Herms & Mattson, 1992; Wahl et al., 2012; Ren et al., 2022). This strategy is consistent with the Optimal Defence Theory, which predicts that chemical defences are preferential allocation of defensive compounds to tissues or regions of highest ecological value (McKey, 1974; Paul et al., 2006). For benthic macroalgae, the thallus surface is continuously exposed to bacterial colonization, biofilm development, epiphytic overgrowth and subsequent macrofouling, making it the most valuable location for chemical defence (Steinberg & de Nys, 2002; Abdul Malik et al., 2020a; Ren et al., 2022; Coolahan & Whalen, 2025).

Together, these findings suggest that *H. floresii* employs a constitutive surface chemical barrier through the selective enrichment of metabolites already present within the whole-cell metabolome, rather than through the production of metabolites exclusive to the thallus surface. Such a strategy enables continuous protection at the organism–seawater interface while avoiding the metabolic costs associated with synthesizing a distinct surface-specific chemical repertoire.

### 4.2 Comparison with chemically defended red macroalgae

The metabolomic profile observed in *H. floresii* agrees with an increasing body of evidence demonstrating that red macroalgae employ chemically mediated defence mechanisms against microbial colonization, epibiosis and herbivory. Among the best-studied examples is *Delisea pulchra*, which produces halogenated furanones that interfere with bacterial quorum sensing and effectively prevent biofilm formation without exhibiting broad-spectrum toxicity (Givskov et al., 1996; Manefield et al., 1999; de Nys et al., 2006; Coolahan & Whalen, 2025). Unlike *Delisea*, whose defence relies on a relatively small family of brominated metabolites, the present study suggests that *H. floresii* may rely on a chemically more diverse mixture dominated by lipid-derived metabolites.

Species belonging to the genus *Laurencia* similarly accumulate halogenated terpenes and acetogenins on the thallus surface, compounds that display antimicrobial, antifouling and antiherbivore activities (Vairappan et al., 2001; Cabrita et al., 2010; Carpena et al., 2023). In contrast, *Asparagopsis taxiformis* synthesizes brominated haloforms and other halogenated metabolites that function as broad-spectrum chemical defences and have recently attracted attention for their antimethanogenic activity (McGurrin et al., 2023). Although brominated compounds have previously been detected in *H. floresii* (Abdul Malik et al., 2020a), the pathway enrichment observed here indicates that lipid metabolism rather than halogenated secondary metabolism dominates the detectable surface metabolome.

The ecological role of lipid-derived metabolites has also been demonstrated in *Gracilaria vermiculophylla,* where mechanical damage rapidly induces the production of oxylipins derived from arachidonic acid metabolism (Nylund et al., 2011; Rempt et al., 2012). Those metabolites function as signaling molecules that regulate wound responses and defence against herbivores and microbial pathogens (Barbosa et al., 2016; Jagusch et al., 2020). The predominance of arachidonic acid metabolism identified in the present study therefore places *H. floresii* within an emerging group of red macroalgae in which lipid-derived signaling molecules constitute important components of chemical defence.

Taken together, these comparisons suggest that while chemically defended Rhodophyta employ different classes of metabolites—including halogenated compounds, terpenoids and oxylipins—the common ecological strategy is the localization of bioactive metabolites at the thallus surface, where they mediate interactions with surrounding microorganisms and fouling organisms.

### 4.3 Surface enrichment of lipid-associated putative metabolites as an energetically efficient surface defence strategy

One of the most notable findings of this study was the predominance of lipid-associated putatives among those preferentially enriched in the surface-associated (DIP) extracts. These metabolites—including fatty acids, oxylipin-like compounds, sterol precursors, sphingolipids and phospholipid intermediates—indicate that lipid metabolism is a major component of the surface chemical defence system of *H. floresii*. Although these metabolites were also present in the whole-cell metabolome, their marked enrichment at the thallus surface suggests that ecological function is determined by spatial allocation rather than exclusive biosynthesis.

The preferential accumulation of lipid-associated metabolites may also represent an energetically efficient defence strategy. Compared with polar metabolites, lipophilic compounds exhibit lower solubility in seawater and are therefore expected to diffuse more slowly from the algal surface (Abdul Malik et al., 2020a; Pozas-Schare et al., 2025). Consequently, their retention within the extracellular mucilage, cell wall matrix or plasma membrane could prolong chemical protection while reducing continual metabolite loss and the energetic costs associated with replenishment Abdul Malik et al., 2020b; Lang et al., 2024). Although this mechanism remains to be experimentally verified in *H. floresii*, it provides a plausible explanation for the dominance of lipid-derived metabolites in the surface extracts. Consistent with this interpretation, experimental studies on the chemically defended red alga *Bonnemaisonia hamifera* have demonstrated that, despite the metabolic cost of chemical defence, its substantial benefits in preventing bacterial colonization and tissue damage favour energetically efficient strategies that maximize protection while minimizing metabolic investment (Nylund et al., 2013; Semmouri et al., 2024).

This hypothesis is further supported by observations from other chemically defended red algae, where defensive metabolites are selectively concentrated at the thallus surface. In *Delisea pulchra*, brominated furanones accumulate at the algal surface, where they disrupt bacterial quorum sensing and inhibit biofilm formation (Dworjanyn et al., 1999; Manefield et al., 1999; Mazumder et al., 2024). Likewise, *Bonnemaisonia hamifera* localizes brominated secondary metabolites in specialized gland cells, maintaining elevated surface concentrations that effectively reduce bacterial colonization (Nylund et al., 2008; Persson et al., 2011). Collectively, these studies support the concept that spatial localization, rather than metabolite novelty, is a key determinant of chemical defence efficacy.

Taken together, these findings suggest that the chemical defence of *H. floresii* is optimized not only through the production of bioactive metabolites but also through their selective enrichment at the thallus surface. We propose that the retention of relatively hydrophobic lipid-associated metabolites at the organism–seawater interface represents an evolutionarily advantageous strategy that prolongs defensive activity, minimizes metabolite loss through diffusion, and reduces the energetic costs of maintaining constitutive surface protection.

### 4.4 Linking metabolomics to the QSI study

A key outcome of the present metabolomic analysis is that it provides a biochemical basis for the quorum-sensing inhibitory (QSI) activity previously reported for surface extracts of *H. floresii* (Abdul Malik et al., 2020a). Using the same surface extracts which inhibited quorum sensing in several Gram-negative bacteria naturally associated with the algal surface, including *Vibrio, Tenacibaculum, Maribacter* and *Aquimarina*. Because quorum sensing regulates bacterial communication, biofilm formation and the establishment of fouling communities, these observations suggested that *H. floresii* employs an effective chemically mediated antifouling strategy.

The present study demonstrates that these biologically active surface extracts are predominantly enriched in lipid-associated metabolites, particularly fatty acids, oxylipin-like compounds and sterol-related metabolites. Fatty acids and oxylipins are increasingly recognized as signalling molecules capable of disrupting bacterial communication, suppressing biofilm development and modulating microbial community composition (Barbosa et al., 2016; Ren et al., 2022; Parchemin et al., 2023). Rather than implicating a single bioactive metabolite, our findings support the hypothesis that the QSI activity of *H. floresii* arises from the collective action of a chemically complex, lipid-rich surface metabolome acts synergistically to suppress microbial colonization, thereby reducing the establishment of both microfouling and subsequent macrofouling communities.

Together, these findings support a coherent ecological model in which *H. floresii* selectively enriches lipid-associated metabolites at the thallus surface, where they interfere with bacterial signalling during the earliest stages of surface colonization. By preventing the establishment of bacterial biofilms, this constitutive chemical barrier is likely to reduce subsequent recruitment of macrofouling organisms, providing an effective first line of defence at the organism–seawater interface.

### 4.5. GC–MS analysis corroborates the lipid-dominated surface metabolome

Complementary GC–MS analysis provided independent validation of the untargeted LC–MS metabolomic findings by directly identifying lipid-derived compounds, including saturated and unsaturated fatty acids, arachidonic acid derivatives, long-chain hydrocarbons and diterpenoid alcohols (see Supplementary Section, Table S6). While LC–MS enabled broad metabolome coverage and pathway-level reconstruction, GC–MS confirmed the predominance of lipid-associated metabolites within the surface extracts, strengthening the conclusion that lipid metabolism is a defining feature of the surface chemistry of *H. floresii*. Previous studies investigating the seasonal variation in the fatty acid composition of *H. floresii* using GC–MS identified three saturated fatty acids (SAFAs; C14:0, C16:0, and C18:0) and one monounsaturated fatty acid (MUFA; C18:1 Δ9). These results indicated a relatively constant SAFA/MUFA ratio throughout the year, except during the windy season, when MUFA levels increased, particularly those of oleic acid (Godínez Ortega et al., 2012).

Several of the identified compounds have established ecological functions in marine macroalgae. Fatty acids and their oxygenated derivatives participate in antimicrobial defence, oxidative stress responses and chemical signalling (Desbois & Smith, 2010; Goecke et al., 2010), whereas phytol and neophytadiene, products of chlorophyll degradation and terpenoid metabolism, exhibit antioxidant and antimicrobial activities (de Morais et al., 2015; Islam et al., 2018). Notably, the detection of arachidonic acid derivatives and eicosapentaenoic acid methyl ester complement the LC–MS pathway analysis by providing direct chemical evidence for the involvement of arachidonic acid metabolism and its associated oxylipin biosynthetic pathways, which are widely implicated in defence and stress signalling in red macroalgae (Bouarab et al., 2004; Ritter et al., 2008; Nylund et al., 2011). The occurrence of these metabolites further supports the hypothesis that surface-associated lipids contribute not only to membrane structure but also to ecological interactions occurring at the algal–seawater interface.

Particularly noteworthy is the identification of arachidonic acid derivatives and eicosapentaenoic acid methyl ester. These polyunsaturated fatty acids serve as precursors of oxylipins, which function as signalling molecules regulating wound responses, defence against herbivores and microbial interactions in numerous red macroalgae (Bouarab et al., 2004; Ritter et al., 2008; Nylund et al., 2011). Their occurrence therefore provides experimental support for the LC–MS pathway analysis, which identified arachidonic acid metabolism as one of the predominant pathways represented among the surface-enriched metabolites.

The GC–MS data also reinforce the compelling evidence of the predominance of lipophilic metabolites enabling prolonged retention at the surface likely adapted for persistent ecological defence. Rather than representing an independent line of evidence, the GC–MS data serve as an orthogonal validation of the untargeted metabolomic analysis. The strong agreement between the two analytical platforms confirms that the surface metabolome of *H. floresii* is dominated by lipid-associated metabolites.

### 4.6. Efficiency and effectiveness of the metabolomic workflow for investigating low-abundance surface metabolites

The characterization of surface-associated metabolites in marine macroalgae remains analytically challenging because of the small quantities of material recoverable from the thallus surface. In *H. floresii*, the limited yield of surface extracts restricted the feasibility of conventional compound isolation and structural elucidation. To address this constraint, we adopted an integrated metabolomic strategy combining untargeted LC–MS profiling, statistical feature selection, putative metabolite annotation, pathway mapping and complementary GC–MS analysis. This approach enabled the chemical complexity of low-abundance surface extracts to be examined at the metabolite-class and pathway levels, providing a basis for ecological interpretation without relying on the definitive identification of individual compounds.

An important strength of this strategy was the convergence of evidence from complementary analytical and computational approaches. Annotation using xMSannotator, supported by cross-referencing against chemical databases and pathway mapping with the Lilikoi R package, consistently indicated the enrichment of lipid-associated metabolic features in surface extracts. GC–MS analysis independently revealed a predominance of fatty acid derivatives and other lipophilic constituents, lending further support to the broader chemical patterns inferred from LC–MS. However, these findings must be interpreted within the limitations of untargeted metabolomics. Most metabolite assignments remain putative, and differences in compound coverage and ionization or detection characteristics between analytical platforms prevent direct confirmation of individual identities. Accordingly, the principal value of the integrated workflow lies in its ability to identify coherent biochemical patterns from chemically limited samples, while providing a foundation for subsequent targeted structural and functional investigations.

By integrating complementary analytical platforms with robust computational interpretation, the approach transformed a chemically sparse dataset into a coherent model of constitutive surface defence. Beyond *H. floresii*, this workflow provides a practical framework for investigating surface metabolomes and chemically mediated interactions in other marine macroalgae, particularly where sample quantity constrains conventional natural-product discovery.

## 5. Study Limitations

Metabolite annotation was largely based on high-confidence putative identification rather than confirmation using authentic standards for all compounds. These limitations are well recognized in untargeted metabolomics studies (Sumner et al., 2007; Uppal et al., 2017; Fiehn, 2002). Consequently, metabolite identities should be interpreted as putative annotations (MSI Level 4), and the biological interpretation is more robust at the level of metabolite classes and metabolic pathways than at the level of individual compounds.

## 6. Conclusion and Future Perspectives

This study presents the first untargeted metabolomic profile of the surface-associated metabolome of *Halymenia floresii*, and it shows that lipid-associated metabolic features are preferentially enriched at the surface extracts compared with whole-cell. Rather than indicating a chemically distinct set of surface-specific compounds, the findings suggest that selective enrichment of metabolites already present within the thallus may be an important feature of the alga’s surface chemistry. This offers new insight into how marine macroalgae might maintain constitutive chemical defence while minimizing the energetic costs associated with metabolite production and loss. In the context of previously reported quorum-sensing inhibitory activity of *H. floresii* surface extracts, this pattern is consistent with a constitutive chemical defence strategy in which lipid-associated metabolites may contribute to maintaining a protective chemical environment at the thallus–seawater interface. Such enrichment could potentially enhance local metabolite availability while limiting losses to the surrounding seawater, although the mechanisms of surface retention and their energetic implications remain to be established experimentally.

Future work should determine which metabolites contribute to these putative defensive functions and how their distribution is regulated. High-resolution tandem mass spectrometry, molecular networking and comparison with authentic standards will be key for increasing confidence in metabolite identification, while spatially resolved mass spectrometry could show their localization across the thallus surface. Combining these methods with bioassay-guided fractionation, quorum-sensing inhibition assays and bacterial colonization experiments would help link metabolite enrichment directly to ecological function. Ultimately, clarifying these links will provide us a more mechanistic understanding of constitutive chemical defence in *H. floresii* and how surface-associated metabolites shape interactions between marine macroalgae and their microbial communities.

## 7. Declarations

### Data availability

Raw data for LC-ESI (+)/MS experiments were deposited and are publicly available in the MetaboLights platform (will be updated after journal submission). GC-MS raw data are available on request from the authors. The authors confirm that the data supporting the findings of this study are available within the article and its supplementary materials.

### Author Contributions

The following statements should be used “conceptualization, S.A.A.M, and N.B.; methodology, S.A.A.M and G.B; formal analysis, S.A.A.M; investigation, N.B, and D.R.; writing—original draft preparation, S.A.A.M; writing—review and editing, N.B.; D.R. and G.B. supervision, Y.F.P.”.

## Supporting information

Supplementary Tables_Abdul Malik et al_Untargeted metabolomics reveals a selectively enriched surface metabolome in the red alga Halymenia floresii

## Acknowledgments

The authors thank L. Taupin for her skillful technical assistance for chromatographic analyses. IBT. V. Avila for culturing the *Halymenia floresii* material used in this experiment.

## References

A Abdul Malik, S, Bedoux, G, Garcia Maldonado J.Q, Freile-Pelegrín Y, Robledo, D & Bourgougnon N (2019) Defence on surface: macroalgae and their surface-associated microbiome. Adv Bot Res. 10.1016/bs.abr.2019.11.009

A Abdul Malik, S, Bedoux, G., Robledo, D., García-Maldonado, J. Q., Freile-Pelegrín, Y., & Bourgougnon, N. (2020b). Chemical defense against microfouling by allelopathic active metabolites of Halymenia floresii (Rhodophyta). J Appl Phycol., 32, 1–15. 10.1007/s10811-019-01999-z

Al-Akwaa, F. M., Yunits, B., Huang, S., Alhajaji, H., & Garmire, L. X. (2018). Lilikoi: an R package for personalized pathway-based classification modeling using metabolomics data. GigaScience, 7(12), giy136. 10.1093/gigascience/giy136

Barbosa, M., Valentão, P. & Andrade, P.B. (2016). Biologically Active Oxylipins from Enzymatic and Nonenzymatic Routes in Macroalgae. Mar Drugs. 14(1):23. doi: 10.3390/md14010023.

Blunt, J. W., Carroll, A. R., Copp, B. R., Davis, R.A., Keyzers, R.A. & Prinsep, M. R. (2018). Marine Natural Products. Nat Prod Rep., 35, 8–53. 10.1039/c7np00052a.

Bouarab, K., Adas, F., Gaquerel, E., Kloareg, B., Salaün, J.P. & Potin, P. (2004) The innate immunity of a marine red alga involves oxylipins from both the eicosanoid and octadecanoid pathways. Plant Physiol. 135(3):1838–48. doi: 10.1104/pp.103.037622.

Cabrita, M. T., Vale, C., & Rauter, A. P. (2010). Halogenated Compounds from Marine Algae. Mar Drugs, 8(8), 2301–2317. 10.3390/md8082301.

Carpena, M., Garcia-Perez, P., Garcia-Oliveira, P. Chamorro, F., Otero, P., Lourenço-Lopes, C., H Cao, Simal-Gandar, J. & Prieto, M.A. (2023). Biological properties and potential of compounds extracted from red seaweeds. Phytochem Rev 22, 1509–1540. 10.1007/s11101-022-09826-z.

Carpena, M., Silva, A., Chamorro, F., Echave, J., Jorge, A. O. S., Barroso, M. F., & Prieto, M. A. (2024). Exploring algal metabolism: insights from metabolomics and computational approaches. Biol Life Sci Forum, 35(1), 9. 10.3390/blsf2024035009.

Coolahan, M. & Whalen, K.E. (2025). A review of quorum-sensing and its role in mediating interkingdom interactions in the ocean. Commun Biol 8, 179. 10.1038/s42003-025-07608-9.

Cronin, G., & Hay, M. E. (1996). Within-plant variation in seaweed palatability and chemical defenses: Optimal Defence Theory versus the Growth-Differentiation Balance Hypothesis. Oecologia, 105, 361–368. 10.1007/BF00328739.

Desbois, A. P., & Smith, V. J. (2010). Antibacterial free fatty acids: activities, mechanisms of action and biotechnological potential. Appl Microbiol Biotechnol., 85, 1629–1642. doi:10.1007/s00253-009-2355-3.

de Morais M.G., Vaz Bda S., de Morais E.G. & Costa J.A. (2015). Biologically Active Metabolites Synthesized by Microalgae. Biomed Res Int. 835761. doi:10.1155/2015/835761.

de Nys, R., Dworjanyn, S. A., Steinberg, P. D. (1998). A new method for determining surface concentrations of marine natural products on seaweeds. Mar Ecol Prog Ser., 162, 79–87.

de Nys, R., Givskov, M., Kumar, N., Kjelleberg, S., & Steinberg, P. D. (2006). Furanones. In N. Fusetani & A. S. Clare (Eds.), Antifouling Compounds (Progress in Molecular and Subcellular Biology, Vol. 42, pp. 55–86). Springer, Berlin. 10.1007/3-540-30016-3_2.

Dworjanyn, S. A., de Nys, R., & Steinberg, P. D. (1999). Localisation and surface quantification of secondary metabolites in the red alga *Delisea pulchra*. Mar Biol., 133, 727–736. 10.1007/s002270050514

Egan, S., Harder, T., Burke, C., Burke, C., Steinberg, P., Kjelleberg, S. & Thomas, T. (2014). The seaweed holobiont: understanding seaweed–bacteria interactions. FEMS Microbiol Rev., 38, 462–476. 10.1111/1574-6976.12011.

Fahy, E., Subramaniam, S., Murphy, R.C., Nishijima, M., Raetz, C.R.H., Shimizu, T., Spener, F., van Meer, G., Wakelam, M.J.O., & Dennis, E.A. (2009). Update of the LIPID MAPS comprehensive classification system for lipids. J Lipid Res., 50 (Suppl), S9–S14. 10.1194/jlr.R800095-JLR200.

Fernie, A. R., Trethewey, R. N., Krotzky, A. J., & Willmitzer, L. (2004). Metabolite profiling: from diagnostics to systems biology. Nat Rev Mol Cell Biol., 5, 763–769. 10.1038/nrm1451.

Fiehn, O. (2002). Metabolomics—the link between genotypes and phenotypes. Plant Mol Biol., 48, 155–171. 10.1023/A:1013713905833.

Gaubert, J., Greff, S., Thomas, O.P., & Payri, C.E. (2019a). Metabolomic variability of four macroalgal species of the genus *Lobophora* using diverse approaches. Phytochem., 162, 165–172. 10.1016/j.phytochem.2019.03.002.

Gaubert, J., Payri, C.E., Vieira, C., Solanki, H., & Thomas, O.P. (2019b). High metabolic variation for seaweeds in response to environmental changes: A case study of the brown algae *Lobophora* in coral reefs. Scientific Rep., 9, 993. 10.1038/s41598-018-38177-z.

Givskov, M., de Nys, R., Manefield, M., Gram, L., Maximilien, R., Eberl, L., Molin, S., Steinberg, P. D., & Kjelleberg, S. (1996). Eukaryotic interference with homoserine lactone-mediated prokaryotic signalling. J Bacteriol., 178(22), 6618–6622. 10.1128/jb.178.22.6618-6622.

Godínez-Ortega, J.L., Ramírez, D.R., Freile-Pelegrín, Y., & Robledo, D. (2012). Seasonal fatty acid composition of *Halymenia floresii* (Rhodophyta) from Yucatán, Mexico. Revista Latinoamericana de Química, 40, 99–105.

Goecke, F., Labes, A., Wiese, J., & Imhoff, J. F. (2010). Chemical interactions between marine macroalgae and bacteria. Mar Ecol Prog Ser, 409, 267–299. 10.3354/meps08607.

Harlin, M. M., & Rice, E. L. (1987). Allelochemistry in marine macroalgae. Crit. Rev. Plant Sci., 5, 237–249. 10.1080/07352688709382241.

Holdt, S. L., & Kraan, S. (2011). Bioactive compounds in seaweed: functional food applications and legislation. J Appl Phycol., 23, 543–597. 10.1007/s10811-010-9632-5.

Horai, H., Arita, M., Kanaya, S., Nihei, Y., Ikeda, T., Suwa, K., Ojima, Y., Tanaka, K., et al. (2010). MassBank: A public repository for sharing mass spectral data for life sciences. J Mass Spectrom., 45, 703–714. 10.1002/jms.1777.

Islam, M. T., Ali, E. S., Uddin, S. J., Shaw, S., Islam, M. A., Ahmed, M. I., Shill, M. C., Karmakar, U. K., Yarla, N. S., et al. (2018). Phytol: A review of biomedical activities. Food Chem Toxicol., 121, 82–94. 10.1016/j.fct.2018.08.032.

Jagusch, H., Baumeister, T. U. H., & Pohnert, G. (2020). Mammalian-like inflammatory and pro-resolving oxylipins in marine algae. Chem bio chem, 21(17), 2419–2424. 10.1002/cbic.202.

Johnson, C. H., Ivanisevic, J., & Siuzdak, G. (2016). Metabolomics: beyond biomarkers and towards mechanisms. Nat Rev Mol Cell Biol., 17, 451–459. 10.1038/nrm.2016.25.

Kanehisa, M., Furumichi, M., Sato, Y., Kawashima, M., & Ishiguro-Watanabe, M. (2023). KEGG for taxonomy-based analysis of pathways and genomes. Nucleic Acids Research, 51, D587–D592. 10.1093/nar/gkac963.

Kumar, M., Kuzhiumparambil, U., Pernice, M., Jiang, Z., & Ralph, P.J. (2016). Metabolomics: An emerging frontier of systems biology in marine macrophytes. Algal Res., 16, 76–92. 10.1016/j.algal.2016.02.033.

Lang, T., Cummins, S. F., Paul, N. A., & Campbell, A. H. (2024). Molecular responses of seaweeds to biotic interactions: A systematic review. J Phycol., 60, 1036–1057. 10.1111/jpy.13504.

Manefield, M., de Nys, R., Naresh, K., Roger, R., Givskov, M., Peter, S. & Kjelleberg S. (1999). Evidence that halogenated furanones from *Delisea pulchra* inhibit acylated homoserine lactone (AHL)-mediated gene expression by displacing the AHL signal from its receptor protein. Microbiol., 145(2):283–91. 10.1099/13500872-145-2-283.

Maximilien, R., de Nys, R., Holmström, C., Gram, L., Givskov, M., Crass, K., Kjelleberg, S., & Steinberg, P. D. (1998). Chemical mediation of bacterial surface colonisation by secondary metabolites from the red alga *Delisea pulchra*. Aquat Microb Ecol., 15, 233–246. 10.3354/ame015233.

Mazumder, S., Bhattacharya, D., Nag, M. & Lahiri, D. (2024). Bioactive compounds from marine algae and fungi in down-regulating quorum sensing. Blue Biotechnology 1, 16. 10.1186/s44315-024-00018-2.

McCall, A. C., & Fordyce, J. A. (2010). Can optimal defence theory be used to predict the distribution of plant chemical defences? J Ecol., 98, 985–992. 10.1111/j.1365-2745.2010.01693.

McKey, D. (1974). Adaptive patterns in alkaloid physiology. The American Naturalist, 108, 305–320. 10.1086/282909.

McGurrin, A., Maguire, J., Tiwari, B. K., & Garcia-Vaquero, M. (2023). Anti-methanogenic potential of seaweeds and seaweed-derived compounds in ruminant feed: current perspectives, risks and future prospects. J Anim Sci. Biotechnol., 14(1), 145. 10.1186/s40104-023-00946-w.

Meesala, S., Gurung, P., Karmodiya, K., Subrayan, P., & Watve, M.G. (2018). Isolation and structure elucidation of halymeniaol, a new antimalarial sterol derivative from the red alga *Halymenia floresii*. J Asian Nat Prod Res., 20, 391–398.

Meldau, S., Erb, M., & Baldwin, I. T. (2012). Defence on demand: mechanisms behind optimal defence patterns. Ann Bot., 110(8), 1503–1514. 10.1093/aob/mcs212.

Nylund, G. M., Cervin, G., Persson, F., Hermansson, M., Steinbery, P.D. & Pavia, H. (2008). Seaweed defence against bacteria: a polybrominated 2-heptanone from the red alga *Bonnemaisonia hamifera* inhibits bacterial colonisation. Mar Ecol Prog Ser., 369, 39–50. 10.3354/meps07577.

Nylund, G. M., Weinberger, F., Rempt, M., & Pohnert, G. (2011). Metabolomic assessment of induced and activated chemical defence in the invasive red alga *Gracilaria vermiculophylla*. PLoS ONE, 6(12), e29359. 10.1371/journal.pone.0029359.

Nylund, G. M., Enge, S., & Pavia, H. (2013). Costs and benefits of chemical defence in the red alga *Bonnemaisonia hamifera*. PLoS ONE, 8, e61291. 10.1371/journal.pone.0061291.

Parchemin, C., Raviglione, D., Mejait, A., Sasal, P., Faliex, E., Clerissi, C. & Tapissier-Bontemps, N. (2023) Antibacterial activities and life cycle stages of *Asparagopsis armata*: Implications of the metabolome and microbiome. Mar. Drugs, 21, 363. 10.3390/md21060363.

Park, E., Yu, H., Lim, J. H., Hee Choi, J., Park, K. J., & Lee, J. (2023). Seaweed metabolomics: A review on its nutrients, bioactive compounds and changes in climate change. Food Res Int. (Ottawa, Ont.), 163, 112221. 10.1016/j.foodres.2022.112221.

Patti, G. J., Yanes, O., & Siuzdak, G. (2012). Metabolomics: the apogee of the omics trilogy. Nat Rev Mol Cell Biol., 13, 263–269. 10.1038/nrm3314.

Paul, N. A., de Nys, R., & Steinberg, P. D. (2006). Chemical defence against bacteria in the red alga *Asparagopsis armata*: linking structure with function. Mar Ecol Prog Ser, 306, 87–101.

Paul, V. J., & Puglisi, M. P. (2004). Chemical mediation of interactions among marine organisms. Nat Prod Rep., 21, 189–209. 10.1039/b302334f.

Paul, N. A., de Nys, R., & Steinberg, P. D. (2006). Seaweed–herbivore interactions at a small scale: Direct tests of feeding deterrence by filamentous algae. Mar Ecol Prog Ser, 323, 1–9. 10.3354/meps323001.

Persson, F., Svensson, R., Nylund, G. M., Fredriksson, N. J., Pavia, H., & Hermansson, M. (2011). Ecological role of a seaweed secondary metabolite for a colonizing bacterial community. Biofouling, 27, 579–588. 10.1080/08927014.2011.589001.

Pozas-Schacre, C., Bischoff, H., Raviglione, D. Chaib, S., Clerissi, C., Bonnard, I., & Maggy, M. N. (2025). Invasive macroalgae shape chemical and microbial waterscapes on coral reefs. Commun Biol 8, 16. 10.1038/s42003-024-07433-6

Qian, P.-Y., Cheng, A., Wang, R., & Zhang, R. (2022). Marine biofilms: Diversity, interactions and biofouling. Nat Rev Microbiol., 20, 671–684. 10.1038/s41579-022-00744-7

Ren, C. G., Liu, Z. Y., Wang, X. L., & Qin, S. (2022). The seaweed holobiont: from microecology to biotechnological applications. Microb biotechnol, 15(3), 738–754. 10.1111/1751-7915.14014

Rempt, M., Weinberger, F., Grosser, K., Pohnert, G. (2012). Conserved and species-specific oxylipin pathways in the wound-activated chemical defense of the noninvasive red alga *Gracilaria chilensis* and the invasive *Gracilaria vermiculophylla*. Beilstein J. Org. Chem., 8, 283–289. 10.3762/bjoc.8.30.

Rhoades, D. F. (1979). Evolution of plant chemical defense against herbivores. In G. A. Rosenthal & Janzen, D. H. (Eds.), Herbivores: their interaction with secondary plant metabolites (pp. 3–54). New York, NY: Academic Press.

Ritter, A., Goulitquer, S., Salaün, J.P., Tonon, T., Correa, J.A., & Potin, P. (2008). Copper stress induces biosynthesis of octadecanoid and eicosanoid oxygenated metabolites in the brown alga *Laminaria digitata*. New Phytol., 180, 809–821. 10.1111/j.1469-8137.2008.02626.x.

Semmouri, I., Janssen,C. R. & Asselman, J. (2024). Allelopathy in macroalgae: Ecological principles, research opportunities and pitfalls reviewed. J Appl Phycol 36 (1): 441–58. 10.1007/s10811-023-03110-z.

Smith, C. A., Want, E. J., O’Maille, G., Abagyan, R., & Siuzdak, G. (2006). XCMS: processing mass spectrometry data for metabolite profiling using nonlinear peak alignment, matching, and identification. Anal Chem., 78(3), 779–787. 10.1021/ac051437y.

Steinberg, P.D. & De Nys, R. (2002). Chemical mediation of colonization of seaweed surfaces. J Phycol., 38: 621–629. 10.1046/j.1529-8817.2002.02042.x.

Steinberg, P.D., de Nys, R. & Kjelleberg, S. (2002) Chemical Cues for Surface Colonization. J Chem Ecol 28, 1935–1951. 10.1023/A:1020789625989

Sumner, L.W., Amberg, A., Barrett, D., et al. (2007). Proposed minimum reporting standards for chemical analysis. Metabolomics, 3, 211–221. 10.1007/s11306-007-0082-2.

Uppal, K., Walker, D. I. & Jones, D. P. (2017). xMSannotator: an R package for network-based annotation of high-resolution metabolomics data. Anal. Chem. 89, 1063–1067. 10.1021/acs.analchem.6b01214.

Vairappan, C. S., Suzuki, M., Abe, T., & Masuda, M. (2001). Halogenated metabolites with antibacterial activity from the Okinawan Laurencia species. Phytochem., 58, 517–523. 10.1016/S0031-9422(01)00260-6.

Wei, R., Wang, J., Su, M. et al. Missing value imputation approach for mass spectrometry-based metabolomics data. Sci Rep 8, 663 (2018). 10.1038/s41598-017-19120-0.

Wahyuni, D.S., Wiryawan, K.G., Ridwan, R., Rafi, M., Gunawan, G., Nurjanah, N., Kusumaningrum, S., et al (2026). Untargeted metabolomic data of some Indonesian seaweed species (*Halymenia durvillei*, *Gracilaria gigas*, Caulerpa racemosa, and Palmaria palmata). Data in Brief, 66, 2352–3409. 10.1016/j.dib.2026.112716

